# Female meiotic drive preferentially segregates derived metacentric chromosomes in *Drosophila*

**DOI:** 10.1101/638684

**Authors:** Nicholas B. Stewart, Yasir H. Ahmed-Braimah, Daniel G. Cerne, Bryant F. McAllister

## Abstract

A vast diversity of karyotypes exists within and between species, yet the mechanisms that shape this diversity are poorly understood. Here we investigate the role of biased meiotic segregation—i.e., meiotic drive—in karyotype evolution. The closely related species, *Drosophila americana* and *D. novamexicana*, provide an ideal system to investigate mechanisms of karyotypic diversification. Since their recent divergence, *D. americana* has evolved two centromeric fusions: one between the 2nd and 3rd chromosomes, and another between the X and 4th chromosomes. The 2-3 fusion is fixed in *D. americana*, but the X-4 fusion is polymorphic and varies in frequency along a latitudinal cline. Here we evaluate the hypothesis that these derived metacentric chromosomes segregate preferentially to the egg nucleus during female meiosis in *D. americana*. Using two different methods, we show that the fused X-4 chromosome is transmitted at an average frequency of ~57%, exceeding expectations of 50:50 Mendelian segregation. Three paracentric inversions are found in the vicinity of the X-4 fusion and could potentially influence chromosome segregation. Using crosses between lines with differing inversion arrangements, we show that the transmission bias persists regardless of inversion status. Transmission rates are also biased in *D. americana*/*D. novamexicana* hybrid females, favoring both the X-4 and 2-3 fused arrangements over their unfused homologs. Our results show that meiotic drive influences chromosome segregation in *D. americana* favoring derived arrangements in its reorganized karyotype. Moreover, the fused centromeres are the facilitators of biased segregation rather than associated chromosomal inversions.

## Introduction

Evolution has produced remarkable diversity in karyotypes among species, but the underlying mechanisms remain unclear. Various mutational events, such as fusion of non-homologous chromosomes, inversions, deletions, duplications or translocation of chromosomal regions are known to alter chromosome number and/or form. Such rearrangements are common but can lead to aneuploidy and/or negative fitness effects that decrease the likelihood of fixation in a population (Bengtsson 1980). One possible mechanism by which these rearrangements can become fixed is centromere-associated meiotic drive (Sandler and Novitski 1957; Pardo-Manuel de Villena and Sapienza 2001a). This hypothesis posits that female meiotic drive shapes karyotype structure through preferential segregation of specific centromere forms. Indeed, in a survey of >1000 mammalian karyotypes, Pardo-Manuel de Villena and Sapienza found that most species either have nearly all metacentric or all acrocentric chromosomes indicating biases that may be shaped by preferential meiotic segregation (Pardo-Manuel de Villena and Sapienza 2001b).

For meiotic drive to operate during gametogenesis, three requirements must be fulfilled: (1) asymmetric meiosis that produces less than four gametes from the four meiotic products, (2) karyotypically heterozygous homologs that pair during meiosis, and (3) asymmetry in the spindle attachment strength between meiotic poles (Sandler and Novitski 1957; Pardo-Manuel de Villena and Sapienza 2001a). Female meiosis is asymmetric because it produces a single functional product; one of four meiotic products forms the functional egg nucleus and the remaining three form the polar bodies. Thus, nonrandom segregation of a given chromosomal variant to either of these two fates indicates the presence of female meiotic drive.

Nonrandom segregation during meiosis can be caused by differences in the strength and/or number of meiotic spindles deployed by the opposing centrioles during metaphase I, or by distinct characteristics of centromere sequences and centromere-associated proteins (Henikoff *et al.* 2001; Chmátal *et al.* 2017). This mechanistic model is supported by recent evidence in mice (Chmátal *et al.* 2014), where the strength of centromeres is determined by the levels of kinetochore proteins localized at the centromere. These proteins are the centromeric attachment site for spindle microtubules, and because different centromeric forms can contain varying levels of these proteins, meiotic drive can favor alternate centromere forms (Chmátal *et al.* 2014).

Two closely related sister-species in the *Drosophila virilis* species group, *D. americana* and *D. novamexicana*, present an excellent opportunity to study the influence of female meiotic drive on chromosome evolution. All members of the *D. virilis* group, except *D. americana*, maintain the ancestral *Drosophila* karyotype of 6 acrocentric chromosomes, or “Muller elements” (Figure 1). In contrast, *D. americana* has evolved two different chromosomal fusions that join the centromeres of non-homologous chromosomes (Figure 1). One centromeric fusion is between the 2nd and 3rd chromosomes (Muller elements D and E), whereas the other is between the X and 4th chromosomes (Muller elements A and B). The X-4 fusion exhibits a latitudinal frequency cline throughout its range in North America (McAllister 2002; McAllister *et al.* 2008): northern populations show a high frequency of the X-4 fusion. Natural selection appears to maintain the alternate chromosome forms across the species range, where the X-4 fusion is favored in cooler, northern latitudes. This study investigates the existence of an intrinsically biased transmission rate favoring the derived fusions present in *D. americana*.

**Figure 1.**
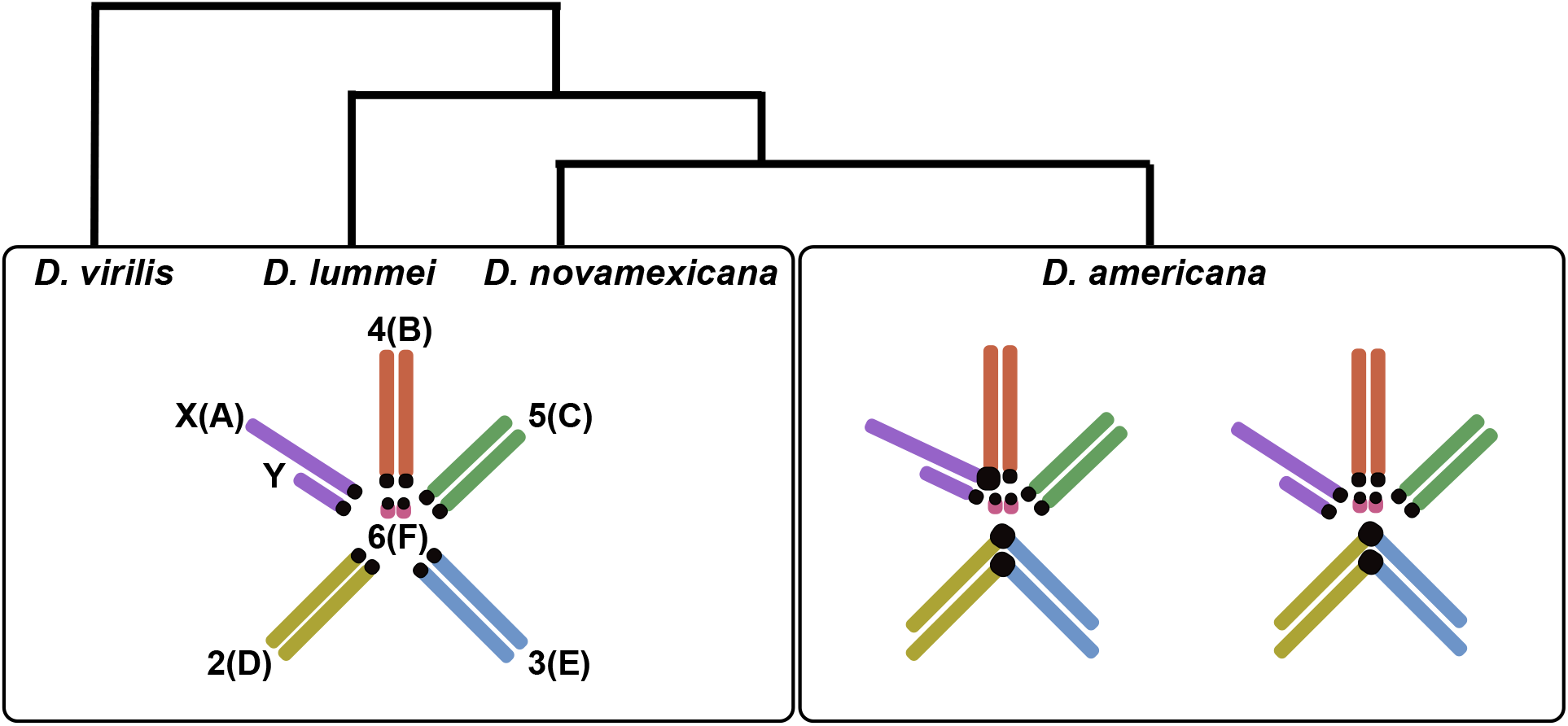
*D. virilis*, *D. lummei*, and *D. novamexicana* have the ancestral karyotype, which consists of five acrocentric, rod-shaped chromosomes and a single “dot” chromosome. *D. americana* has evolved two centromeric fusions between chromosomes 2 and 3 (fixed) and chromosomes X and 4 (polymorphic). The Muller element classification of each chromosome is indicated in parentheses.

Chromosomal inversions can potentially influence chromosome segregation by influencing the formation of dicentric or acentric chromatids following crossing over. For example, during female meiosis, a meiocyte has the ability to pull dicentric or acentric chromatids—which arise from a cross-over event within a heterozygous inverted region—towards the polar body during meiosis. This can ensure that the egg nucleus will inherit the functional monocentric chromatid (Sturtevant and Beadle 1936; Carson 1946). This mechanism could potentially affect segregation of chromosomes that differ in inversion content and produce meiotic drive. In *D. americana*, multiple paracentric inversions reside near the X-4 centromere. Two different inversions are observed together on the 4th chromosome of *D. americana*: the smaller *In(4)b* inversion is always found nested within the larger *In(4)a* inversion. However, not every fused X-4 chromosome has this inversion complex (Hsu 1952; Evans *et al.* 2007). This inversion complex is also not present within the unfused 4th chromosome of *D. americana*. In contrast, *D. novamexicana* is fixed for the *In(4)a* inversion and lacks the *In(4)b* inversion. A third inversion, *In(X)c*, is present on the X chromosome and is perfectly associated with the X-4 fusion but is not found on the unfused arrangement of *D. americana* (Hsu 1952). It is, however, present on the unfused X chromosome of *D. novamexicana*.

Here we analyze the transmission dynamics of the derived metacentric chromosomes of *D. americana*, both in within-species heterozygotes and in *D. americana*/*D. novamexicana* hybrids. We use two methods to track the transmission of the two chromosomal types. First, we use a visible eye color mutation on the 4th chromosome to track the inheritance of the X-4 fusion from heterozygous females to adult sons. Second, we use microsatellite markers that are tightly associated with the centromeres to track the inheritance of the X-4 and 2-3 fusions from heterozygous females to early-stage embryos. These two approaches allow us to assess the impact of differential viability on transmission ratios. Finally, we analyze transmission rates of several combinations of inversion heterozygotes to assess the effects of the three associated inversions on the transmission of the X-4 fusion. Our results show that biased meiotic transmission favors the derived, fused arrangements present in *D. americana* at an average rate of ~57%. This bias favors the X-4 fusion regardless of inversion status, suggesting that centromeres are likely the causal factor influencing meiotic drive. Furthermore, we find no difference between our assessment of meiotic drive in embryos and adults, indicating that differential viability is not an important factor in the observed transmission bias.

## Materials and Methods

### Measuring transmission of the fused X-4 chromosome using a visible marker

The transmission rate of the fused and unfused chromosomes was measured in F1 females obtained in reciprocal crosses between five *D. americana* strains with the X-4 fusion (SB02.02, SB02.06, SB02.08, SB02.10, *red*) and five strains with the unfused arrangement (ML97.3, ML97.4, ML97.5, ML97.6, *pur*). (See Table S1 for description of strains). In all crosses, virgin flies were collected within 36 hours of eclosion and kept at 22°C until sexually mature at ~7 days. We performed 45 out of 50 possible reciprocal combinations of ‘fused by unfused’ crosses. To generate heterozygous females, four virgin adult females from a fused (or unfused) strain were crossed with four virgin adult males from an unfused (or fused) strain. Approximately 30 crosses of individual F1 females mated with *cardinal* (*cd*) males that carry a recessive red eye color mutation on the 4th chromosome were established to enable segregation of the fused and unfused arrangements. F2 male sires were backcrossed to *cd* females, and their F3 progeny were evaluated for an association between eye color and sex (Figure 2). F2 males inherit the fused X-4 chromosome if all F3 males show the *cd* phenotype and all females are wild-type. On the other hand, F2 males inherit the unfused X and 4th chromosomes if the *cd* phenotype is present in equal proportions to the wild-type in both F3 males and F3 females. We measured the F1 female transmission ratio from ≥100 F2 males for each heterozygous genotype combination.

**Figure 2.**
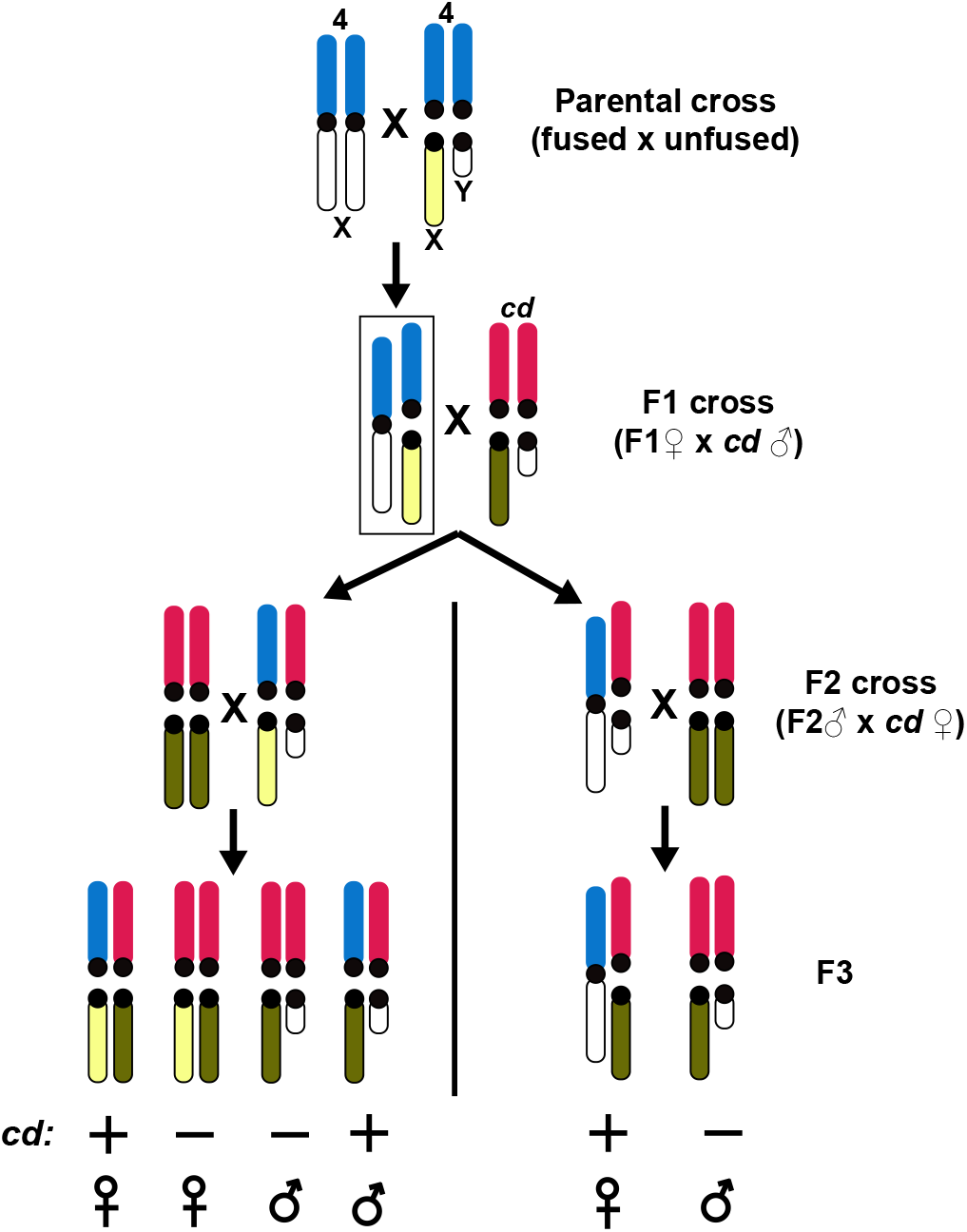
Crossing scheme used to track the transmission of the X-4 fused and unfused arrangements from F1 heterozygous females (rectangle) to F2 adult sons. Parental 4th chromosomes are shown in blue, and the *cd*-containing 4th chromosome is shown in red. The fused X, unfused X, and *cd*-derived X chromosomes are shown in white, yellow, and green, respectively.

### Measuring transmission of the fused X-4 chromosome using microsatellite markers

#### Microsatellite marker design

Microsatellite markers were developed near the centromeres of the X, 4th, and 3rd chromosomes to track the transmission of the X-4 and 2-3 fused chromosomes from heterozygous females to their offspring. Descriptions of twenty markers near the centromere of the X-4 fusion and nine markers near the centromere of chromosome 3 are provided in Table S2. The three *D. virilis* genomic scaffolds we used for marker development were mapped previously to the centromere-proximal euchromatin of *D. virilis* polytene chromosomes by Schaeffer *et al*. (2008). We identified 20-50bp tandem repeats in the *D. virilis* genome sequence using RepeatMasker (Smit *et al.* 2013). We chose repeats that were closest to the proximal end of the scaffold (closest to the centromere) for further investigation. We designed primer pairs that flank the repeat regions using Primer3 (Untergasser *et al.* 2012). Primer sequences were blasted against the draft assembly of *D. americana* (Reis *et al.* 2008) to check for sequence conservation and multiple annealing sites. We checked for allelic differences among ten inbred *D. americana* and four *D. novamexicana* strains. We ultimately obtained five informative microsatellite markers that can be used to track transmission of the X-4 fusion, and two informative markers to track transmission of the 2-3 fusion (Table S2).

#### Embryo collection

Two different heterozygous female genotypes were used in this experiment. The first genotype was generated by crossing two inbred *D. americana* lines (fused X-4: G96.23, unfused: HI99.12), and the second, interspecific genotype was generated from a cross between an inbred *D. americana* strain (G96.13) and an iso-female *D. novamexicana* strain (1031.0). Parental crosses were performed as described above. For the intraspecific F1 cross, ~100 heterozygous F1 females were crossed with males from a different unfused inbred *D. americana* strain (ML97.5). For the interspecific F1 cross, ~100 heterozygous F1 females were crossed with males from a different *D. novamexicana* strain (1031.4). Females were allowed to mate and lay eggs in 12-hour intervals on grape agar plates, followed by an additional 10-12 hour incubation at 22°C to allow embryos to develop. Embryos were collected daily over an 8-day period to ensure sampling from the females’ entire sexual peak. Collected embryos were rinsed with distilled water, washed with 50% bleach, and frozen at −20°C in 96-well plates. DNA was prepared from frozen embryos using a standard squashing buffer (10mM Tris-HCl pH 8.2, 1mM EDTA, 25mM NaCl, 200μl/ml Proteinase K).

#### Microsatellite analysis

Each microsatellite locus was amplified from an embryonic DNA sample using a previously developed PCR method (Shimizu *et al.* 2002). This method entails three PCR primers at differing concentrations: 26.7nM of a modified forward primer, 133.3nM of an IR-700 or IR-800 conjugated M13 forward primers, and 160nM of the reverse primer. The forward primer was synthesized with an addition of the reverse complement of an M13 tail (5’-GGATAACAATTTCACACAGG) at the 5’ end. This modification introduces an M13 complement on the PCR amplicon, which is used to produce dye labeled single-strand products from either the IR-700 or IR-800 conjugated M13 primer (LI-COR, Lincoln, NE). Genotypes were determined following separation on a 7% polyacrylamide gel and imaged using the LI-COR Odyssey infrared imager with Odyssey application software v3.0 (Figure 3).

**Figure 3.**
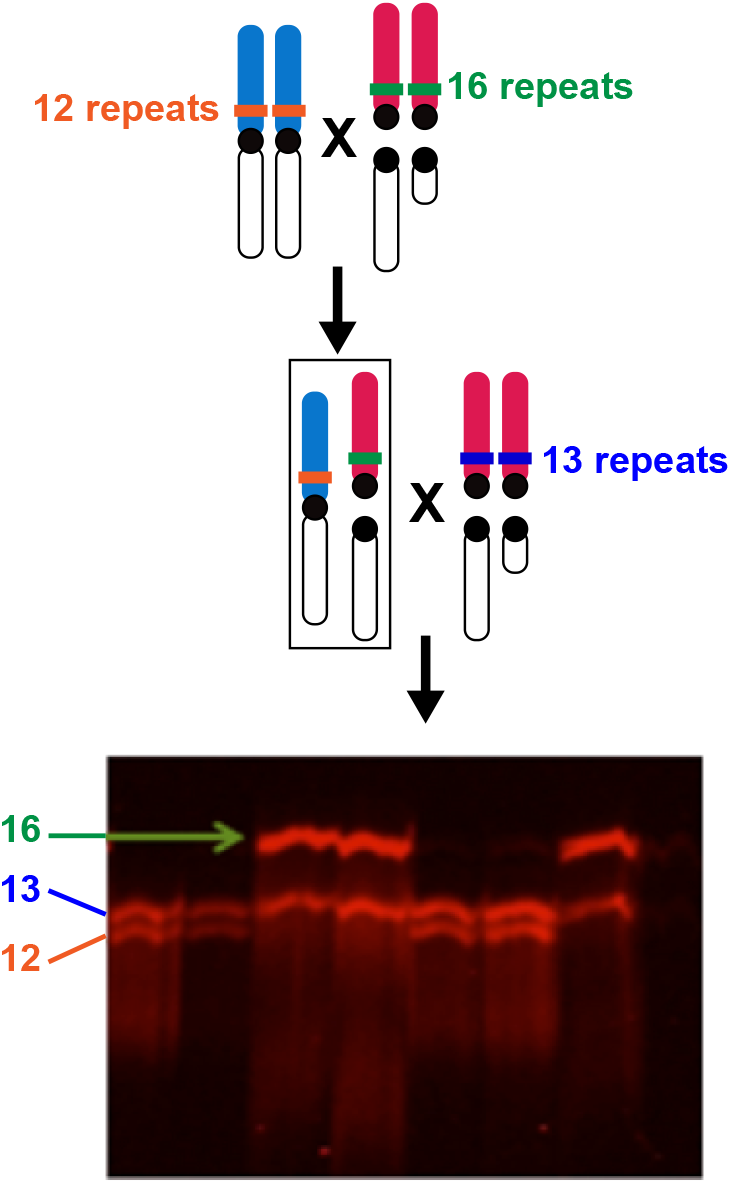
Crossing scheme to track transmission of the X-4 fusion from F1 females (rectangle) by microsatellite genotyping of three alleles in F2 embryos.

We determined the transmission of the fused X-4 chromosome from F1 G96.23/HI99.12 females by genotyping embryos at the microsatellite locus of the 4th chromosome, ms1019560 (Table S2). Inbred lines G96.23, HI99.12, and ML97.5 are each homozygous for different length alleles at ms1019560. The F2 offspring each have the band corresponding to the paternal ML97.5 line and another band that either corresponds to the G96.23 X-4 chromosome or the HI99.12 unfused 4th chromosome (Figure 3). Transmission rates of the X-4 fused chromosome were calculated as the number of F2 embryos with the G96.23 genotype divided by the total number of embryos genotyped.

To determine the transmission rate of the X-4 fused chromosome in interspecific F1 hybrid females (G96.13/1031.0), F2 embryos were genotyped using three separate microsatellite loci on the 4th chromosome: ms977861, ms1219821, and ms2825734 (Table S2). At each locus, G96.13 and 1031.0 had distinct microsatellite alleles. However, the strain that hybrid F1 females were crossed with (1031.4) had the same allele at all three loci. Therefore, the F2 embryos from this cross were screened for the G96.13 allele. Offspring that inherit the fused X-4 chromosome will carry both the 1031.0/1031.4 and G96.13 alleles, but those that inherit the unfused chromosome will only carry the 1031.0/1031.4 allele.

### Evaluating the effect of the chromosomal inversions in transmission ratio distortion

Inbred lines with previously confirmed inversion rearrangements (Mena 2009) were used to analyze the effects of the different inversions on transmission bias. Two inbred fused X-4 lines that have *In(4)ab* and *In(X)c* were used: G96.13 and HI99.34 (Table S1). Another two lines, G96.23 and OR01.50, have the In(X)c inversion. The unfused X and 4th arrangement in *D. americana* lacks all three inversions associated with X-4 fusion (inbred lines HI99.12 and *pur*). The two *D. novamexicana* lines contain both *In(4a)* and *In(X)c*, but lack *In(4)b*.

These lines were systematically crossed to generate three female genotypes that are heterozygous for the X-4 fusion, but also contain different heterozygous combinations of the three inversions: (1) Females that are heterozygous for all three inversions were generated by crossing G96.13 females with *pur* males (Figure 4A), (2) females heterozygous for *In(X)c* we generated by crossing G96.23 or OR01.50 females with HI99.12 males (Figure 4B), and (3) females heterozygous for *In(4)b* but homozygous for the other two inversions were generated by crossing G96.13 or HI99.34 females with males from the *D. novamexicana* strains, 1031.0 or 1031.4, respectively (Figure 4C). Transmission of the fused X-4 chromosome was tracked using the *cd* visible eye marker, as outlined above and in Figure 2.

**Figure 4.**
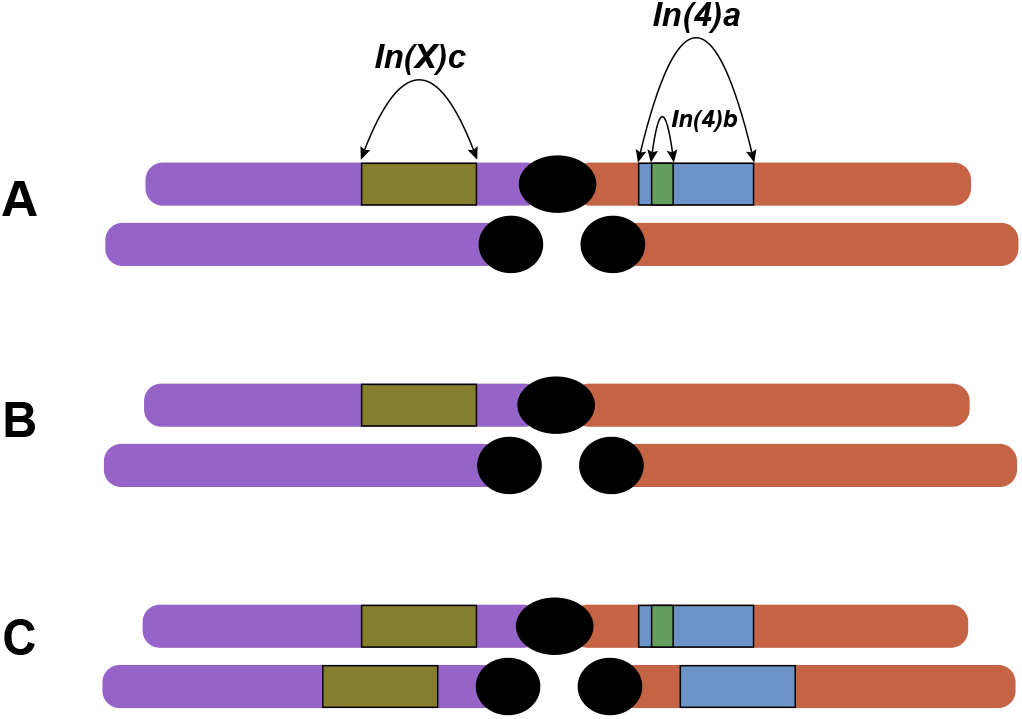
Visual representation of F1 females with the three different possible inversion states. A) Heterozygous for all three inversions. B) X-4 fused chromosome is lacking In(4)ab, therefore only In(X)c is heterozygous. Transmission ratio of the fused X-4 chromosome can be measured without the effects of In(4)ab. C) D. americana/D. novamexicana hybrid, In(4)a and In(X)c are homozygous and only In(4)b is heterozygous. Transmission ratio of the fused X-4 chromosome can be measured without the effects of In(X)c.

### Assaying transmission of the fused 2-3 chromosome of D. americana

We tested whether meiotic drive causes transmission bias of another fused metacentric chromosome, chromosome 2-3, by measuring transmission rates from *D. americana*/*D. novamexicana* hybrid females to adult offspring. Heterozygous females were generated by crossing strain G96.13 (*D. ame*) with 1031.4 (*D. nov*), which were subsequently backcrossed to 1031.4 and her progeny frozen for DNA extraction. Whole fly DNA was prepared as described previously, and samples were analyzed at two microsatellite loci (ms786503 and ms806741; Table S2) on the 3rd chromosome. F2 offspring were screened for the presence of the G96.13 allele, which would indicate transmission of the fused 2-3 chromosome from the heterozygous F1 female. F2’s from this cross were also genotyped at two additional loci on the X and 4th chromosomes (ms1141205 and ms1219821, respectively; Table S2).

### Statistical analysis

We used a logistic generalized linear model with maximum likelihood fitting to test for significant deviations from Mendelian expectations of X-4 transmission (File S1). The model was used to examine whether maternal or paternal inheritance of the X-4 fusion affected the transmission rate, and to obtain transmission rate estimates and 95% confidence intervals for each cross (File S1). Analyses comparing the transmission rates to embryos and adults from specific lines were tested against Mendelian expectations using a χ^2^ goodness-of-fit test: A 2×2 contingency table with χ^2^ was used to compare crosses to each other. The effects of different inversion rearrangements were analyzed using a χ^2^ goodness-of-fit test against the Mendelian expectation of 50:50, and a 2×2 contingency table with a χ^2^ test was used to compare inversion rearrangements against each other.

## Results

### The X-4 fusion exhibits a transmission advantage over its unfused homolog in D. americana

We performed reciprocal crosses between five parental unfused X and 4 lines and five parental fused X-4 lines to produce F1 females that are heterozygous for the alternate arrangements of the X and 4th chromosomes. Transmission rates from heterozygous F1 mothers to sons were measured by tracking sex-linkage of a phenotypic marker on chromosome 4 (Figure 2). An average transmission rate of 56.6% (95% *C. I.* = 1.8%) for the X-4 fusion was observed among the F1 females produced from 45 different cross combinations. Introducing the X-4 fusion maternally or paternally does not significantly influence the transmission rate (Figure 5A, Table S3, File S1); including this factor in a logistic generalized linear model did not improve the fit of the model (χ^2^=2.4, *p*=0.12, *d.f.*=1, File S1).

**Figure 5.**
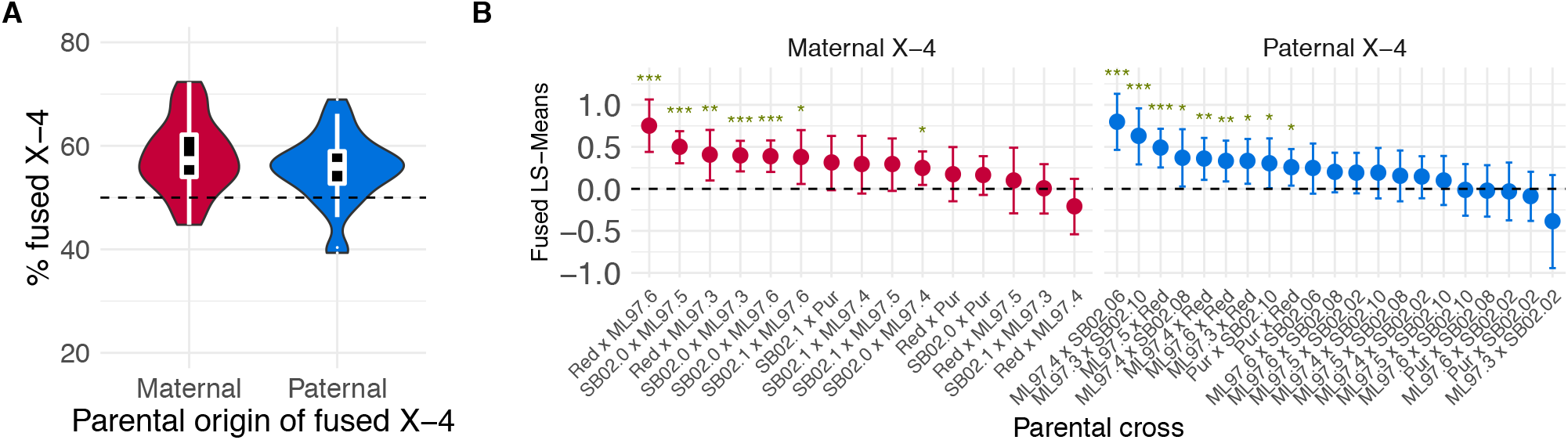
Transmission ratio of the fused X-4. (A) Females heterozygous for the arrangement of the X and 4th chromosomes were produced from pairwise reciprocal crosses between lines differing in chromosome arrangement, and transmission ratios are presented for females that inherited the X-4 fusion maternally or paternally. (B) Least squares means estimates of transmission ratios for the fused X-4 chromosome for each cross combination. The dotted line (LS-Means = 0) indicates 50:50 Mendelian segregation, and positive values indicate biased transmission favoring the X-4 fusion. Error bars represent 95% confidence intervals. Cross combinations with significant deviation from Mendelian segregation are indicated (*: 0.01 < *p* < 0.05; **: 0.001 < *p* < 0.01; ***: *p* < 0.001).

Meiotic segregation of the alternative chromosome arrangements in these F1 females is expected to have a 50:50 transmission ratio. The estimated 95% confidence intervals overlapped with the 50:50 Mendelian expectation for 28 out of the 45 heterozygotes produced from inter-strain crosses (Figure 5B, File S1). In contrast, the confidence intervals exceed 50:50 for the remaining 16 genotypes, including instances where the X-4 chromosome was introduced maternally and paternally. In none of the F1 genotypes examined was the estimated confidence interval for the transmission ratio below 50:50 (Figure 5B, File S1). Overall, these results show that the derived metacentric X-4 fusion experiences a transmission bias over the ancestral acrocentric arrangement.

### Meiotic drive is not caused by differential viability

To assess the effect of differential viability on meiotic drive, we measured transmission rates of the X-4 fusion from heterozygous females to adult sons and to embryonic offspring in two separate crosses. We generated intraspecific (G96.23 x HI99.12) and interspecific hybrid (G96.13 x 1031.0) F1 females, and subsequently genotyped their adult male progeny and embryo offspring at a molecular marker on the 4th chromosome. Two separate trials of embryos were analyzed to ensure consistency of the method.

Offspring from the conspecific F1 females showed a transmission bias for the fused X-4 chromosome in both embryos and adult sons (Figure 6). One embryo trial showed a bias of 54.5% and the other a bias of 58.9%; both statistically different from the predicted 50:50 (Table S4). These two embryo trials were not significantly different from each other with respect to transmission bias for the fused X-4 arrangement (χ^2^= 1.03, *p*>0.05, *d.f.*=1), reflecting consistency in replicate trials. Transmission of the X-4 chromosome to adult sons (61.9%) is not significantly different from either embryo replicate (χ^2^= 3.8, *p*>0.05, *d.f.*=1; χ^2^= 0.46, *p*>0.05, *d.f.*=1) (Table S4).

**Figure 6.**
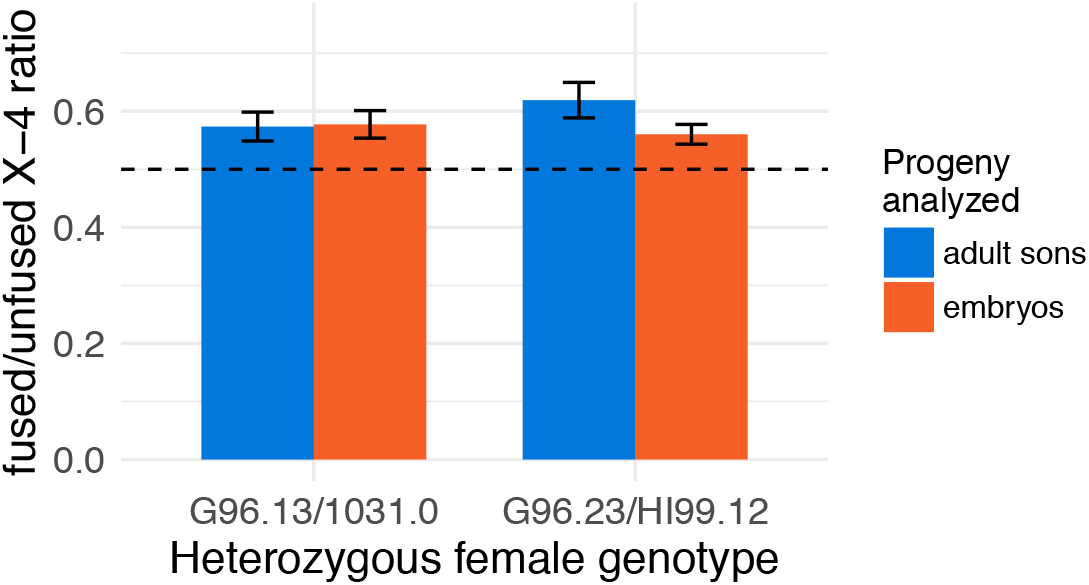
Transmission ratio of the X-4 fused chromosome from conspecific (right) and heterospecific (left) heterozygous females to adult sons and embryos. Error bars represent the binomial standard error.

The assessment of transmission ratio was consistent between embryos and adult sons; however, this consistency was attained with only 77% and 73% success genotyping individual embryos in the two trials. The remaining embryos failed to produce amplified DNA at the microsatellite locus. To assess whether our ability to successfully genotype an embryo was skewing the results in favor of the fused X-4 arrangement, we crossed the parental unfused X and 4th line, HI99.12, to another unfused line, ML97.5. F1 embryos were analyzed to assess the success rate for genotyping the unfused 4th chromosome: if the unfused arrangement is less likely to produce useable DNA for genotyping, it is expected that the embryos from the parental unfused line will produce fewer successfully genotyped embryos. We compared our embryo genotyping between offspring of the parental unfused line and both trials of F1 heterozygous females. The genotyping success rate for offspring originating from the parental female was 75.1% (n=405), which was not significantly different in embryo trials of embryos laid by F1 heterozygous females (2×3 contingency table chi squared; χ^2^= 3.77, *p*>0.1, *d.f.*=2). From this comparison we can conclude that this approach does not introduce a genotyping bias against the unfused arrangement in embryos.

F1 hybrids from the *D. americana*/*D. novamexicana* cross (G96.13 x 1031.0) were also assayed for biased transmission of the X-4 fusion in embryos and adult sons by genotyping at three molecular markers near the centromere of chromosome 4 (ms977861, ms1219821, and ms2825734; Table S2). Marker ms977861 is most proximal to the centromere, roughly 1 Mb from the end of the scaffold, while ms1219821 and ms2825734 are ~250kb and ~1.8Mb away from ms977661, respectively.

The transmission rate of the fused X-4 chromosome was ~57% in both adult sons and embryos (Figure 6, Table S4). In this cross there was also no significant difference between the inheritance rates of the fused X-4 to adult sons and embryo offspring (χ^2^=0.01, *p*>0.05, *d.f.*=1). We observe distinct genotypes between the most distal rearrangement (ms2825734) and the other 2 markers in 1.2% of embryos, suggesting low levels of recombination between these loci. Only one recombinant genotype between ms977861 and ms1219821 was observed, suggesting a recombination rate lower than 0.5%. All samples that showed recombinant genotypes at any of the two loci were not included in the analysis.

### Meiotic drive is not affected by centromere-associated inversions

We examined whether three inversions near the X and 4th chromosome centromeres influence transmission of the X-4 fusion. We generated three combinations of heterozygous inversion genotypes on an X-4 fusion heterozygous karyotype (Figure 4), and measured transmission ratios of the X-4 fused chromosome using the crossing scheme described in Figure 2. First, we assayed meiotic drive in females that are heterozygous for all three known inversions on the X and 4th chromosomes: *In(X)c*, *In(4)a*, and *In(4)b* (Figure 4). In these females, the transmission rate of the fused X-4 chromosome was 57.8% (Figure 7, Table S5), which is significantly higher than Mendelian transmission (χ^2^=7.46, *p*<0.01, *d.f.*=1). Second, we examined the effect of a heterozygous *In(X)c* inversion genotype in an otherwise collinear arrangement of the 4th chromosome—i.e. lacking the *In(4)ab* complex (Figure 4B). Here we used two separate conspecific crosses to produce F1 females: G96.23xHI99.12 and OR01.50xHI99.12. Heterozygous F1 females from both crosses showed a transmission bias for the X-4 fusion (G96.23/HI99.12: χ^2^=13.82, *p*<0.001, *d.f.*=1; OR01.50/HI99.12: χ^2^=3.92, *p*<0.05, *d.f.*=1; Figure 7, Table S5). F1 females from the G96.23/HI99.12 cross showed a higher transmission rate of the fused X-4 chromosome (61.9%) than the OR01.50/HI99.12 F1 females (54.9%), but these transmission rates are not significantly different from each other (χ^2^=3.18, *p*>0.05, *d.f.*=1). Furthermore, F1 females from these two crosses showed no significant transmission difference from F1 females that are heterozygous for *In(4)ab* (χ^2^=3.19, *p*>0.05, *d.f.*=2), suggesting the linear arrangement of the 4th chromosome does not affect meiotic drive for the fused X-4 chromosome.

**Figure 7.**
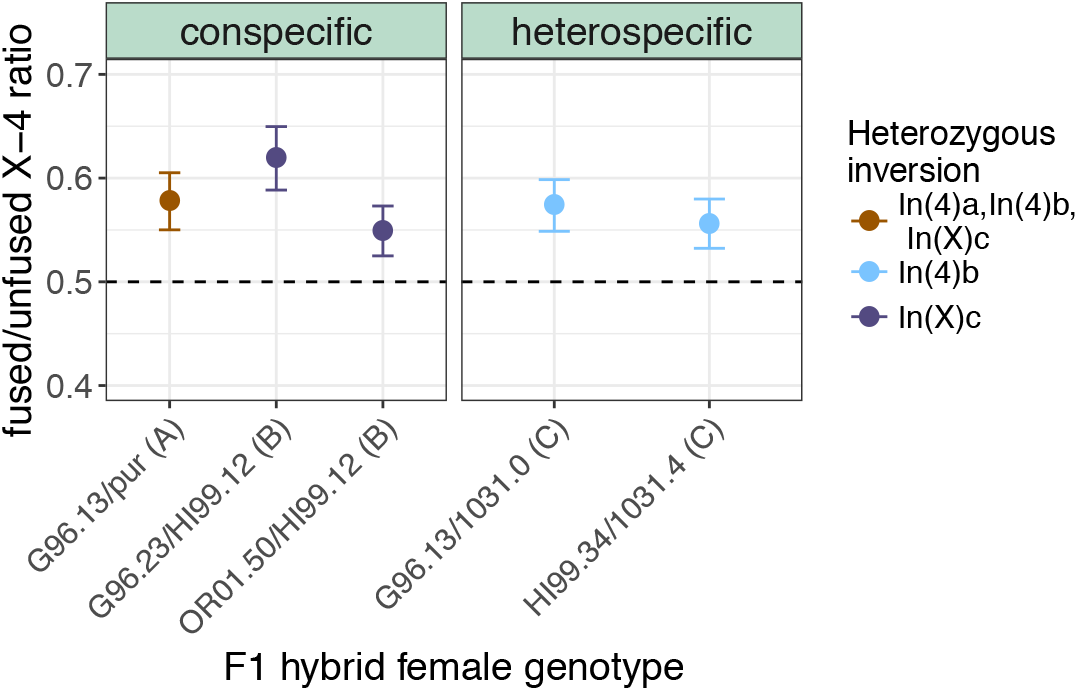
Transmission ratios of the X-4 fusion from heterozygous females to adult sons in crosses with differing inversion states. Point color represents heterozygous inversion genotype (legend on the right), which is also indicated by the letters in parenthesis next to the parental strain identification (see Figure 4). The left and right panels represent con- and heterospecific hybrid females, respectively. Error bars represent the binomial standard error, and a 50:50 segregation ratio is indicated by the dashed line.

Finally, We assessed the effect of the smaller inversion on chromosome 4, *In(4b)*, in an otherwise collinear genotype that is homozygous for *In(X)c* and *In(4)a*. Here, we utilized *D. novamexicana* to generate heterospecific hybrid females, because it also has *In(X)c* but lacks *In(4)b*. We crossed *D. americana* females that had *In(X)c* and *In(4)ab* with *D. novamexicana* males that had *In(X)c* and *In(4)a* (Figure 4C). Two separate crosses (G96.13 x NOVA1031.0 and HI99.34 x NOVA1031.4) were performed to generate F1 females. Again, transmission rates favored the X-4 fused chromosomes (Figure 7). F1 G96.13/NOVA1031.0 females and F1 HI99.34/NOVA1031.4 females had a significant transmission rate of 57.4% and 55.6%, respectively, favoring the X-4 fusion (G96.13/NOVA1031.0: χ^2^=8.24, *p*<0.01, *d.f.*=1; HI99.34/NOVA1031.4: χ^2^=5.28, *p*<0.02, *d.f.*=1; Figure 7, Table S5). These crosses were also not significantly different from each other (χ^2^= 0.26, *p*>0.05, *d.f.*=1). Furthermore, transmission rates did not differ from those in triple inversion heterozygotes (χ^2^=0.426, *p*>0.8, *d.f.*=2).

Taken together, these results indicate that the three centromere-associated inversions on the X and 4th chromosomes play no detectable role in the observed transmission bias that favors the X-4 fusion.

### The fused 2-3 chromosome shows biased transmission in D. americana/D. novamexicana hybrid females

We investigated whether female meiotic drive favors all fused metacentric chromosomes in *D. americana* by measuring the transmission rates of the 2-3 fused chromosome from *D. americana*/*D. novamexicana* hybrid females to adult offspring. F1 hybrid *D. americana*/*D. novamexicana* were produced from a cross between G96.13 females and 1031.4 males, and were heterozygous for both the X-4 and 2-3 fusions. The transmission rate of the 2-3 fused chromosome was determined by microsatellite genotyping of adult offspring at two loci on the 3rd chromosome (ms786503 and ms806741), which are ~200kb apart and show perfect co-segregation. The transmission of the X-4 chromosome was measured by genotyping adult offspring at two loci: one on the X chromosome (ms1141205) and another on the 4th chromosome (ms1219821). Genotypes at these two markers were incongruent less that 1% of the time.

Adult offspring from F1 hybrid females inherited the 2-3 and X-4 fused chromosomes 62.6% and 55.4% of the time, respectively (Table 1). The X-4 transmission rate is similar to the 57% ratio observed in embryo and adult male offspring (χ^2^= 0.72, *p*>0.3, *d.f.*=1). Furthermore, the transmission rate of the 2-3 fused chromosome is significantly greater than the X-4 fused chromosome (χ^2^= 4.29, *p*<0.04, *d.f.*=1).

**Table 1.** Transmission ratio of fused 2-3 and X-4 chromosomes in *D. amer*/*D. nov* hybrid females

| Chr. | # fused | # unfused | Total | % fused | $\chi^2$ | p-value |
| --- | --- | --- | --- | --- | --- | --- |
| 2-3 | 221 | 132 | 353 | 62.6 | 21.9 | 2.87e-06 |
| X-4 | 286 | 231 | 517 | 55.4 | 5.64 | 0.018 |

Finally, we analyzed co-segregation of the fused X-4 chromosome and the fused 2-3 chromosome during meiosis in F1 hybrid females. We genotyped 333 flies for both the X/4th and 2nd/3rd chromosome arrangements. We find no evidence of co-segregation between the X-4 fused chromosome and the 2-3 fused chromosome (χ^2^= 2.2, *p*>0.1, *d.f.*=1) (Table S6). Thus, the forces biasing the transmission for the fused X-4 appear to act independently of the forces driving the fused 2-3 chromosome.

## Discussion

Here we presented evidence of meiotic drive favoring two derived metacentric chromosomes. The fused X-4 chromosome in *D. americana*, which resulted from a fusion between two acrocentric chromosomes, enjoys a transmission advantage of ~53-70% in heterozygous females. In addition, the 2-3 fused chromosome transmission rate is >60% over its unfused counterpart in heterospecific hybrid females. The transmission bias for the fused X-4 chromosome is observed in both 24hr-old embryos and adult sons, providing evidence that the observed meiotic drive is not an artifact of reduced survival of the unfused genotype. Furthermore, the arrangements of the differing inversions do not play a detectable role in the transmission bias, suggesting that the differing centromeres or a locus linked to the centromere are impacting the observed bias. Finally, there is no difference in transmission bias based on the parental origin of the fused X-4 chromosome, suggesting that maternally inherited components are not playing an important role in the transmission bias. We discuss these findings in more detail below.

### Meiotic drive vs. differential viability

We investigated two plausible explanations for the observed transmission bias between the karyotype forms in *D. americana*: meiotic drive and differential viability. Observing the same biased ratio from heterozygous females to embryos and to adult sons suggests that transmission bias takes place during meiosis. However, we were unable to assay every embryo collected due to unsuccessful DNA preparations for some embryos. This is likely due to the observation that some laid eggs are not fertilized, and thus likely contain little DNA for microsatellite amplification. We have shown in previous work that hatch rates within and between *D. americana* strains range from 70%-90%, and are primarily due to lack of fertilization (Ahmed-Braimah and McAllister 2012). However, we found no correlation between fertilization rates and presence/absence of the X-4 fusion in that study.

Another study of fertility/viability and early development between homozygotes of each arrangement at a similar temperature that we maintain the lines in (22°C) showed higher viability for the unfused lines (Sillero *et al.* 2014). This suggests that if differential viability was playing a significant role, the transmission bias would favor the unfused X and 4th arrangement rather than the fused X-4. Without the ability to successfully assay every embryo collected from heterozygous females, we cannot completely rule out the possibility that differential viability in early development between the two chromosome arrangements is involved in biasing the inheritance pattern. However, if this were the case, we would expect the unfused chromosomal arrangement to yield a lower successful embryo genotyping percentage than the embryos of experimental heterozygous F1 females, but this was not observed.

Recent studies have shown that meiotic drive likely contributes to the evolution of chromosomes with differing centromeres such as the yellow monkeyflower, *Mimulus guttatus* (Fishman and Willis 2005; Fishman and Kelly 2015), and the house mouse *Mus musculus* (LeMaire-Adkins and Hunt 2000; Chmátal *et al.* 2014). Meiotic drive has also been hypothesized as a major contributor to mammalian karyotype evolution (Pardo-Manuel de Villena and Sapienza 2001b; Yoshida and Kitano 2012). Taken together, these studies and the findings presented here suggest that meiotic drive is a taxonomically broad and pervasive force impacting karyotype evolution.

### Centromere vs. inversions

We did not observe a significant difference in segregation bias between different inversion states of the fused chromosomes, and all chromosome arrangements showed a bias for fused chromosomes regardless of inversion status. However, currently we are unable to create an F1 female that is homozygous for all three inversions. Thus, it is possible that just the presence of an inversion could cause meiotic drive between the differing centromere arrangements. The most plausible mechanism for this inversion effect would arise when recombination occurs within the inversion break points. This would generate dicentric and acentric chromosomes, which can be selectively eliminated by ensuring transmission of the recombinant chromosome to the polar bodies (Carson 1946). If this were the case, the size of the inversion would be expected to influence the magnitude of the drive as the recombination events would have a greater likelihood of a recombination site inside a larger inversion. Because *In(x)c* is much larger than *In(4)b*, we would expect G96.23/HI99.12 and OR01.50/HI99.12 lines to have greater drive than G96.13/Nova1031.0 and HI99.34/Nova1031.4. Yet, we observe no difference between the lines that only differ at *In(4)b* and the lines that only differ at *In(x)c*, suggesting that the centromeric fusion—or a locus closely linked to the centromere—is the driver of biased segregation. In other systems, meiotic drive is also caused by differing centromere arrangements (Fishman and Saunders 2008) or beta chromosomes that contain centromeric knobs (Buckler *et al.* 1999). Our results strongly implicate the centromere (or centromere-associated sequences) as the target of biased segregation.

### Transmission bias of the 2-3 chromosome vs. X-4 chromosome

We found meiotic drive favoring both the X-4 and the 2-3 chromosomes. Thus, in this system, meiotic drive favors the derived metacentric chromosomes at the expense of the acrocentric homologs. Similar mechanisms likely underlie the transmission bias for the X-4 and 2-3 fused chromosome arrangements. However, the transmission bias for the 2-3 chromosomes appears to be greater than that of the X-4th chromosome. In addition, the 2-3 and the X-4 chromosomes do not co-segregate within the same meiosis. While both metacentric chromosomes are favored in female meiosis, the different magnitude of drive and the lack of co-segregation indicates that the mechanism affecting meiotic drive in *D. americana* are likely complex. Mechanisms of meiotic drive in *M. musculus* are affected by levels of kinetochore proteins surrounding the differing arrangements (Chmátal *et al.* 2014). In populations that have fixed metacentric chromosomes, the relative localization of the kinetochore proteins HEC-1 (Ndc80 in Drosophila) and CENP-A (CID in Drosophila) at metacentric centromeres is significantly higher. In contrast, populations that have all acrocentric chromosome have relatively higher amounts of HEC-1 and CENP-A localizing to acrocentric chromosomes. In mouse populations that have closer to half acrocentric chromosomes and half metacentric chromosomes, variation in relative localization of HEC-1 and CENP-A between acrocentric and metacentric chromosomes is observed (Chmátal *et al.* 2014). Ndc80 in Drosophila (HEC-1) is a component of the ndc80 complex, which is a core component of the kinetochore and is involved in many processes for both mitosis and meiosis, including kinetochore assembly, congression of the chromosomes at the metaphase plate, and binding to the spindle (Tooley and Stukenberg 2011). CID (CENP-A) is the centromere specific histone H3 protein that plays a role in proper kinetochore recruitment and centromere formation (Blower and Karpen 2001). Differences in the observed transmission bias between the 2-3 and the X-4 could result from differing centromere compositions, such that the 2-3 fused chromosome recruits higher levels of kinetochore proteins than the 2nd and 3rd acrocentric chromosomes, and at a more consistent rate than the X-4 over the X and 4th. A difference in centromere composition could be a factor in the observation that the segregation of the X-4 chromosome does not affect segregation of the 2-3 chromosome.

### Control of meiotic drive

Meiotic drive in excess of 50:50 segregation should facilitate driving the favored arrangement to fixation in the absence of an opposing selective force. In nature the X-4 fused arrangement is not fixed in populations, but rather exists in a latitudinal cline throughout the central United States (McAllister 2002; McAllister *et al.* 2008). These contemporary population samples of adult flies from throughout the species range, along with early surveys (Patterson and Stone 1952), reveal the widespread presence and persistence of this chromosomal polymorphism where the derived X-4 fusion is rare in the south and common in the north. The intrinsic segregation advantage for the metacentric chromosome is difficult to reconcile in the context of this apparently stable cline. While stability of the cline has previously been attributed to natural selection in the context of climatic variability, this variability may also affect the mechanism causing meiotic drive. Recombination studies in *Drosophila melanogaster* have suggested that transmission distortion may play a role in increased inheritance of chromatids that have undergone crossing over in females which have been exposed to stress either by bacterial infection or heat treatment (Singh *et al.* 2015; Jackson *et al.* 2015). Moreover, the latitudinal cline may represent a balance between meiotic drive and forces of natural selection acting on the allelic content of the alternative chromosome forms. This would suggest a strong selective advantage for the unfused arrangement in the southern United States which decreases with increasing latitude, until the population is fixed for the fused X-4 chromosome. To further investigate these explanations, transmission rates of heterozygous females exposed to differing stress inducers related to the northern and southern United States should be examined.

