## Supplementary tables for "Female meiotic drive preferentially segregates derived metacentric chromosomes in *Drosophila*"

**Table S1**: Description of strains.

| line name | Species | Type | X-4 Status | Inversions | Collected location | Year | Previously Described |
| --- | --- | --- | --- | --- | --- | --- | --- |
| SB02.02 | *D. americana* | IsoFemale | fused | unknown | Salisbury Rec Area, IA | 2002 | McAllister and Evans 2006 |
| SB02.06 | *D. americana* | IsoFemale | fused | unknown | Salisbury Rec Area, IA | 2002 | McAllister and Evans 2006 |
| SB02.08 | *D. americana* | IsoFemale | fused | unknown | Salisbury Rec Area, IA | 2002 | McAllister and Evans 2006 |
| SB02.10 | *D. americana* | IsoFemale | fused | unknown | Salisbury Rec Area, IA | 2002 | McAllister and Evans 2006 |
| ML97.3 | *D. americana* | IsoFemale | unfused | none | Monroe, LA | 1997 | Mena 2009 |
| ML97.4 | *D. americana* | IsoFemale | unfused | none | Monroe, LA | 1997 | Mena 2009 |
| ML97.5 | *D. americana* | IsoFemale | unfused | none | Monroe, LA | 1997 | Mena 2009 |
| ML97.6 | *D. americana* | IsoFemale | unfused | none | Monroe, LA | 1997 | Mena 2009 |
| Red | *D. americana* | Inbred | fused | *In(4)ab, In(X)c* | Salisbury Rec Area, IA | 2002 | Mena 2009 |
| G96.13 | *D. americana* | Inbred | fused | *In(4)ab, In(X)c* | Gary, IN | 1996 | Mena 2009 |
| G96.23 | *D. americana* | Inbred | fused | *In(X)c* | Gary, IN | 1996 | Mena 2009 |
| OR01.50 | *D. americana* | Inbred | fused | *In(X)c* | Ottawa Refuge, OH | 2001 | Mena 2009 |
| HI99.34 | *D. americana* | Inbred | fused | *In(4)ab, In(X)c* | Howell Island near St. Louis, MO | 1999 | Mena 2009 |
| Pur | *D. americana* | Inbred | unfused | none | Monroe, LA | 1997 | Mena 2009 |
| ML97.5 I-12 | *D. americana* | Inbred | unfused | none | Monroe, LA | 1997 | Mena 2009 |
| HI99.12 | *D. americana* | Inbred | unfused | none | Howell Island near St. Louis, MO | 1999 | Mena 2009 |
| NOVA1031.0 | *D. novamexicana* | Inbred | unfused | *In(4)a, In(X)c* | Grand Junction, CO | before 1984 | UCSD *Drosophila* stock number 15010-1031.04 |
| NOVA1031.4 | *D. novamexicana* | Inbred | unfused | *In(4)a, In(X)c* | Moab, UT | 1949 | UCSD *Drosophila* stock number 15010-1031.04 |

**Table S2:** Microsatellite loci and corresponding primers used in the analyses.

| Locus^1^ | Chrom. | Scaffold | Location^2^ | Annealing temp (˚C) | Forward primer | Reverse primer |
| --- | --- | --- | --- | --- | --- | --- |
| ms399482 | X | 12970 | 399482 | 53 | 5`CCCTTATTTACTTACGTTTGATCG | 5`ATTGGCGTCGATCTGTTTTT |
| ms897939 | X | 12970 | 897939 | 52 | 5`TGAGCGGACTGCCTCTCTAT | 5`CACAAATATGACTTCTCGACCA |
| ms1141205^3^ | X | 12970 | 1141205 | 54 | 5`CTGAACGACAGGCGATTGA | 5`GCAGAGTAATTTCGCCAACC |
| ms1147269 | X | 12970 | 1147269 | 53 | 5`TGCACTTGATAAAAGCCAAGC | 5`TTGCATGTCTTAAGTTGGATGAA |
| ms290883 | 4 | 12723 | 290883 | 54 | 5`CCGGCTAGGTAGATAGGGAGA | 5`AAATCCCTCCGAGGTTTGTTA |
| ms512634 | 4 | 12723 | 512634 | 53 | 5`ACGGCGCATAGCTTTTTATG | 5`CTGTTCTCAAGTAGTTCGCTCTC |
| ms977861^3^ | 4 | 12723 | 977861 | 54 | 5`AATCCAGTACGTCGCCTGTG | 5`GTTCGCCGTCTCTTGCTAAT |
| ms1005794 | 4 | 12723 | 1005794 | 52 | 5`CTCAGAGTTTATCCCAACAGAAA | 5`GCATTTGTCCACATTACTTGAA |
| ms1015661 | 4 | 12723 | 1015661 | 54 | 5`AGCTTGGCTAACGGCACTT | 5`GAGCAAAGCGTGGAAAACTT |
| ms1019560^3^ | 4 | 12723 | 1019560 | 56 | 5`CAAGCGTACTAATTGATCCTG | 5`GCGCCACATTCCTTTCAA |
| ms1219821^3^ | 4 | 12723 | 1219821 | 54 | 5`GTTGCAGTGGCATAGAATTA | 5`TTTAGGTTTTAGGTGGCATT |
| ms1212995 | 4 | 12723 | 1212995 | 55 | 5`TCGCGTATATAAGCCCCATA | 5`GGGAGATTGCCAACTCAAGA |
| ms1221886 | 4 | 12723 | 1221886 | 53 | 5`CTGCTGAGCGTGGGAAAG | 5`AACTGGCCAAATAGCATGAA |
| ms1245189 | 4 | 12723 | 1245189 | 57 | 5`GCAGGGCACTAGTTTTACCC | 5`AGCATCCACTGGAGAACTGG |
| ms1723757 | 4 | 12723 | 1723757 | 53 | 5`CCACTATGTCGCATATGTTCA | 5`ACGAAAAGGACAACAAGGAG |
| ms1962073 | 4 | 12723 | 1965073 | 51 | 5`ATGACTAACGTTGACAGCAC | 5`CCAAACAGAATACACACTG |
| ms2642948 | 4 | 12723 | 2642948 | 50 | 5`GCAACGCAATATGTCAGTTA | 5`CACTCAACATAGGGCAATTA |
| ms2018415 | 4 | 12723 | 2018415 | 51 | 5`TTTAAAGACCACAAAAAGC | 5`CAAAACATGTGTCCCTACTG |
| ms2825734^3^ | 4 | 12723 | 2825734 | 56 | 5`TACGAGCGTATAATGCAAA | 5`ACAAGAAAAAGCTGAAGGAG |
| ms3110818 | 4 | 12723 | 3110818 | 48 | 5`CAGGCGAAAAATACAACTAA | 5`AAATCAACCCTCGAATAATG |
| ms225279 | 3 | 13049 | 24916626 | 55 | 5`TGCAGCCAAAAACAAAACTG | 5`TCAACATCTCTGGCAGCATC |
| ms248590 | 3 | 13049 | 24893315 | 53 | 5`AAATGTTTGCAAGGGTGTGC | 5`TGTACGTATTTTTGTATATGTG |
| ms636836 | 3 | 13049 | 24505069 | 54 | 5`CCATCTCACTTATGGCCTCA | 5`TGTGAAGACGTGCCAAAATG |
| ms724459 | 3 | 13049 | 24417446 | 56 | 5`AAGCTTTTTGGTCTGGCTTG | 5`AACGGCATATAGCTACCACGA |
| ms786503^3^ | 3 | 13049 | 24355402 | 54 | 5`TTCTGTCCGCAAGGGTATT | 5`ACGCGTCTACTGAACCATTC |
| ms806741^3^ | 3 | 13049 | 24335164 | 54 | 5`GCGGTTTTTGGATGTTCACT | 5`TGCACTGAAAAAGTGGCCTA |
| ms827964 | 3 | 13049 | 24313941 | 53 | 5`TTTTGGCTTATATCGTTTTTGGA | 5`AACTGAGTTTTGGACCGATTC |
| ms840661 | 3 | 13049 | 24301244 | 55 | 5`ACACACCCAGCACAACCA | 5`CGACTTACGAATTGTGTCTCCA |
| ms814370 | 3 | 13049 | 24327535 | 54 | 5`GGGCAATACATATTGGTGATGTT | 5`TCCGATAACCGATTACTCATCTC |

^1^ The distance in base pairs from the proximal end of the scaffold is represented given in the marker’s name.

^2^ The location given is the base position within each respective scaffold of the *D. virilis* genome.

^3^ Informative markers used in experiments.

**Table S3:** Transmission ratios of the fused X-4 chromosome from heterozygous females to adult sons.

| Maternal line | Paternal line | Fused | Unfused | % Fused | Total |
| --- | --- | --- | --- | --- | --- |
| Red* | ML 97.3 | 106 | 71 | 59.89 | 177 |
| SB 02.02* | ML 97.3 | 70 | 48 | 59.32 | 118 |
| SB 02.06* | ML 97.3 | 117 | 100 | 53.92 | 217 |
| SB 02.08* | ML 97.3 | 101 | 47 | 68.24 | 148 |
| SB 02.10* | ML 97.3 | 89 | 89 | 50 | 178 |
| Red* | ML 97.4 | 64 | 79 | 44.76 | 143 |
| SB 02.02* | ML 97.4 | 55 | 63 | 46.61 | 118 |
| SB 02.06* | ML 97.4 | 55 | 46 | 54.46 | 101 |
| SB 02.08* | ML 97.4 | 110 | 63 | 63.58 | 173 |
| SB 02.10* | ML 97.4 | 76 | 57 | 66.5 | 133 |
| Red* | ML 97.5 | 53 | 48 | 52.48 | 101 |
| SB 02.02* | ML 97.5 | 88 | 75 | 53.99 | 163 |
| SB 02.06* | ML 97.5 | 117 | 50 | 70.06 | 167 |
| SB 02.08* | ML 97.5 | 74 | 45 | 62.18 | 119 |
| SB 02.10* | ML 97.5 | 92 | 69 | 57.14 | 161 |
| Red* | ML 97.6 | 123 | 58 | 67.96 | 181 |
| SB 02.02* | ML 97.6 | 92 | 74 | 55.42 | 166 |
| SB 02.06* | ML 97.6 | 69 | 42 | 62.16 | 111 |
| SB 02.08* | ML 97.6 | 109 | 67 | 61.93 | 176 |
| SB 02.10* | ML 97.6 | 92 | 63 | 59.35 | 155 |
| Red* | Pur | 81 | 68 | 54.36 | 149 |
| SB 02.02* | Pur | 85 | 73 | 53.8 | 158 |
| SB 02.06* | Pur | 72 | 61 | 54.14 | 133 |
| SB 02.10* | Pur | 87 | 64 | 57.62 | 151 |
| ML 97.3 | Red* | 129 | 93 | 58.11 | 222 |
| ML 97.4 | Red* | 150 | 105 | 58.82 | 255 |
| ML 97.5 | Red* | 190 | 117 | 61.89 | 307 |
| ML 97.6 | Red* | 156 | 112 | 58.21 | 268 |
| Pur | Red* | 186 | 144 | 56.36 | 330 |
| ML 97.3 | SB 02.02* | 21 | 31 | 40.38 | 52 |
| ML 97.4 | SB 02.02* | 132 | 115 | 53.44 | 247 |
| ML 97.5 | SB 02.02* | 146 | 121 | 54.68 | 267 |
| ML 97.6 | SB 02.02* | 64 | 66 | 49.23 | 130 |
| Pur | SB 02.02* | 86 | 94 | 47.78 | 180 |
| ML 97.4 | SB 02.06* | 111 | 50 | 68.94 | 161 |
| ML 97.6 | SB 02.06* | 98 | 77 | 56 | 175 |
| ML 97.4 | SB 02.08* | 81 | 56 | 59.12 | 137 |
| ML 97.5 | SB 02.08* | 91 | 78 | 53.85 | 169 |
| ML 97.6 | SB 02.08* | 153 | 126 | 54.84 | 279 |
| Pur | SB 02.08* | 81 | 83 | 49.39 | 164 |
| ML 97.3 | SB 02.10* | 99 | 53 | 65.13 | 152 |
| ML 97.4 | SB 02.10* | 94 | 78 | 54.65 | 172 |
| ML 97.5 | SB 02.10* | 95 | 86 | 52.49 | 181 |
| ML 97.6 | SB 02.10* | 81 | 82 | 49.69 | 163 |
| Pur | SB 02.10* | 103 | 76 | 57.54 | 179 |

* Strains with fused X-4 chromosomes

**Table S4:** Transmission rates of the X-4 fusion from heterozygous females to adult sons and embryos.

| Offspring | cross | Total | # Fused | # Unfused | % Fused | *χ*^2^ | p value |
| --- | --- | --- | --- | --- | --- | --- | --- |
| adult sons | G96.23/HI99.12 | 252 | 156 | 96 | 61.9 | 13.82 | <0.001 |
| embryos | G96.23/HI99.12 | 568 | 310 | 258 | 54.5 | 4.58 | <.0.05 |
| embryos | G96.23/HI99.12 | 278 | 164 | 114 | 58.9 | 8.23 | <0.01 |
| adult sons | G96.13/1031.0 | 394 | 226 | 168 | 57.4 | 8.24 | <0.01 |
| embryos | G96.13/1031.0 | 433 | 250 | 183 | 57.7 | 10.06 | <0.01 |

**Table S5:** Transmission rates of the X-4 fusion from heterozygous females with different inversion arrangements to adult sons.

| Cross | Heterozygous inversions | total | # fused | # unfused | % fused | *χ*^2^ | p-value |
| --- | --- | --- | --- | --- | --- | --- | --- |
| G96.13/pur | *In(4)a, In(4)b, In(X)c* | 322 | 186 | 136 | 57.75 | 7.46 | <0.01 |
| G96.23/HI99.12 | *In(X)c* | 252 | 156 | 96 | 61.9 | 13.82 | <0.001 |
| OR01.50/HI99.12 | *In(X)c* | 428 | 235 | 193 | 54.9 | 3.92 | <0.05 |
| G96.13/1031.0 | *In(4)b* | 394 | 226 | 168 | 57.4 | 8.24 | <0.01 |
| HI99.341031.4 | *In(4)b* | 437 | 243 | 194 | 55.6 | 5.28 | <0.02 |

**Table S6:** Number of individuals inheriting the four possible X-4 and 2-3 arrangement combinations.

|  | Fused 2-3 | Unfused 2^nd^ and 3rd |
| --- | --- | --- |
| Fused X-4 | 109 | 77 |
| Unfused X and 4th | 99 | 49 |
